# Multi-layer encapsulation of plant protoplasts

**DOI:** 10.1101/2025.05.08.652860

**Authors:** Matthew S. Grasso, Rachael Floreani, Philip M. Lintilhac

**Affiliations:** Department of Plant Biology, The University of Vermont, Burlington, Vermont, USA; Departmentt of Mechanical Engineering, The University of Vermont, Burlington, Vermont, USA

## Abstract

We report here on an attempt to create an engineered structure which can reproduce the physical and mechanical environment of the land plant sporangium. This work is part of a broader effort to understand the stress mechanics of the land plant sporangium and its role in initiating reproductive development. A second purpose is to extend the range of experimental methods available for the study of the physical environment of plant cell growth and the mechanics of trans-cellular signaling in plant development. We describe an experimental protocol, based on the microfluidic encapsulation of living plant protoplasts in multilayered microbeads composed of tunable hydrogel materials whose mechanical properties can be modified to mimic the stress-mechanics of the living sporangium. Our results demonstrate the successful encapsulation of living plant protoplasts in dual-layered microbeads consisting of an inner layer of gelled agarose and an outer layer of cross-linked alginate/methacrylate.

## Introduction

In the animal kingdom eggs and sperm differentiate early in life and are maintained as distinct lineages that persist until sexual maturity. This aspect of sexual development allows for the harvesting of naïve eggs and sperm for in vitro fertilization. But in the land plants there is no comparable lineage of reproductive cells maintained during vegetative growth that is dedicated to gametogenesis and reproduction. Instead, reproductive cells differentiate *de novo* as the plant enters its reproductive phase. The *de nov*o origin of the reproductive germ lines in plants raises fundamental questions about mechanism underlying the shift in developmental fate from mitotic growth to reproductive growth and gametogenesis.

It has been proposed that the land plant sporangium evolved as a stress-mechanical focusing device that targets a precisely located region in the growing sporangium (Lintilhac, 2024), shifting the developmental fate of the most centrally located cells (the archesporium) towards reproductive differentiation and gametogenesis. In this work we take first steps towards reconstructing the physical architecture of the sporangium in an engineered structure. Previous work has shown that living plant protoplasts can be successfully encapsulated in hydrogel microspheres and maintained in vitro for extended periods of time (Grasso & Lintilhac, 2016). Here we introduce a more advanced embedment methodology, showing that we can build multi-layered hydrogel microbeads comprising an inner agarose layer and an outer alginate-methacrylate (AlgMA) layer, suggesting that with further development it may become possible to construct biomimetic sporangial structures that are capable of initiating archesporial differentiation and meiosis *in vitro*.

### Experimental approaches

While the goal of in vitro gametogenesis in plants is unattainable at present, the work reported here describes an experimental protocol designed to physically constrain individual plant cells in ways that may allow us to manipulate and interpret stress, strain, and shear at the level of the individual cell while maintaining the ability to visualize cellular responses at the sub-cellular level.

Plant cells and tissues are equipped with mechanosensitive structures capable of reacting rapidly and precisely to changing load conditions (Schaefer et al 2017) (Mirabet et al, 2018) (Livanos & Muller, 2019). The cellular structures that most likely fill this need are an assemblage of cytoplasmic proteins collectively referred to as the cytoskeleton, consisting of semi-rigid compressive rods known as microtubules, and tensile contractile fibers called microfilaments. The cytoskeleton, augmented by a number of accessory proteins, can together be regarded as a unifying tensegrity structure (Ingber, 2003, Ingber, 2008) capable of reading stress, strain, and shear information at any location in the cytoplasm (Colin, 2020).

The underlying hypothesis is that in the pre-meiotic sporangium, the specification and subsequent channeling of developmental fate towards reproductive differentiation and meiosis depends upon deterministic physical information networks that have overcome many of the limitations of stochastic molecular information systems (Hemenway & Gehring, 2023) (Kurusu, et al, 2013) (Lintilhac, 2022).

Further advances in active hydrogel engineering should make it possible to control the shrinkage and/or swelling of the individual layers (Sugaya, et al, 2013), to create localized isotropic singularities capable of transmitting unique physical cues to individual cells at precise locations in a larger cell mass.

## Materials and Methods

### Terminology

#### Protoplast

living plant cells from which the cellulosic cell wall has been removed.

#### Microdroplet

protoplasts encapsulated in un-gelled agarose droplets.

#### Microsphere

microdroplets that have been cooled to their gelling temperature.

#### Microbeads

microspheres that have been re-encapsulated in a second layer of crosslinked hydrogel (alginate-methacrylate).

### Hydrogel Polymers

1. **Agarose:** Sigma-Aldrich Inc. USA, low gelling temperature Agarose #A9045.
2. **(AlgMA) Alginate Methacrylate:** Lyophilized AlgMA polymer was provided by the Floreani Lab in the Dept. of Mechanical Engineering, University of Vermont, USA. (Tahir and. Floreani, 2022).

### Solutions and Culture Media

1. **(CWDS) Cell Wall Digestion Solution:** 9% mannitol, 0.05% MgCl2, 0.4% Cellulase RS, 0.4% Cellulase R-10, 0.05% Macerozyme R-10, 0.05% Pectolyase Y-23, 20 microliters Rohapect UF, 20 microliters Rohapect 10L, 20 microliters protease inhibitor cocktail, 0.19% MES and pH adjusted to 5.8 using KOH. The solution was sterilized using a rapid flow Nalgene filter unit 0.43µm.
2. **(PWS #1) Protoplast Washing Solution:** 9.3% mannitol, 0.05% MgCl_2_, and 0.1% MES, pH to 5.8.
3. **(PWS#2) Protoplast Washing Solution w/o MgCl**_**2:**_ 9.3% mannitol, 0.1% MES pH to 5.8.
4. **Conditioned Medium** was collected from actively growing, 4-day old *Nicotiana tabacum* cv. BY-2 suspension cultures by filtration through a rapid flow 0.43-micron filter. Conditioned medium was stored in a freezer without autoclaving.
5. **(PCM) Protoplast Culture Medium:** 3% sucrose, 0.43% MS Medium (Caisson Labs), 0.2mg/L 2,4-dichlorophenoxyacetic acid, 1mg/L thiamine, 100mg/L myo-inositol, 4.5% mannitol, 10mL Conditioned Medium, and pH adjusted to 5.8 using 0.1 M KOH.
6. **(CBS) Cell Buoyancy Solution:** 0.5% sucrose, 1.65% high MW Dextran 150000, 8.65% mannitol, 0.1% MES, 0.04% MgCl_2_, 4% (v/v) OptiPrep™, pH adjusted to.
7. **(CS) Calcium Solution** 4% mannitol, 1.46% CaCl_2_ 2H_2_O, pH to 5.8 using 0.1M KOH.
8. **(AMDS) Alginate Methacrylate Droplet Solution** 7.5% mannitol, 1% F68 Pluronic, and 0.073% CaCl_2_ 2H_2_O (5mM CaCl_2_), pH to 5.8 using 0.1 M KOH.

*All prepared media were autoclave sterilized unless otherwise noted.

### Step 1

Protoplast isolation.

Although many sources of living plant protoplasts can be proposed, we began with an established plant cell culture line that is immortal and easily cultured to prepare wall-less protoplasts in quantity.

*Nicotiana tabacum* BY-2 cell line (rpc00041) was provided by RIKEN BRC through the National BioResource Project of the MEXT, Japan. Protoplasts were obtained from 4-day suspension cultures grown at 27°C in MS Medium (Murashige and Skoog basal salts, 3% sucrose, 100 mg/L myo-inositol, 1 mg/L thiamine, and 0.2 mg/L 2,4-dichlorophenoxyacetic acid).

Suspension-cultured cells were removed from the culture medium and enzymatically stripped of their cell walls in cell wall digestion solution (CWDS). Suspension cultured BY-2 cells were digested for 2-4 h at 27×C on a shaker table. The time required to generate viable protoplasts from suspension culture material varied depending on the cell line, the batch of enzyme solution, and the starting cell material, therefore the digestion was split into two parts. During the first part BY-2 cells in digestion medium were placed in an incubating shaker (27° C, 120 R.P.M.) for between 45 Minutes and 1 hour 20 Minutes. The first part of the digestion was stopped when most cells had become spherical in shape but were still attached to their neighbors. They were then transferred to an incubator (27° C, no shaking) for between 1hour 15 minutes and 2 hours 40 minutes. The second step of the protoplast generation process was stopped when almost all protoplasts had been released from their neighbors. Protoplasts were separated from the digestion medium by centrifugation at 100g for 7 Minutes and washed twice in protoplast wash solution 1 (PWS #1). Protoplasts were then resuspended in a cell buoyancy solution (CBS) and diluted to a concentration of 2250 protoplasts/µL in preparation for the encapsulation process.

### Step 2

Single layer microsphere production.

Agarose microspheres were generated using a commercial microfluidic droplet system (Dolomite Microfluidics). Water-based (agarose) droplets were formed as the discontinuous phase in a 2-reagent, 4-channel, hydrophobic junction chip # 3200242 (Junction diameter 100u). The continuous phase was a light mineral oil (Sigma Aldrich M5310) with 2% Span 80 (Fluka Analytical, St. Louis, Missouri, USA) added to prevent droplet coalescence. Flow rates of the protoplasts and agarose were adjusted with separate Dolomite P-Pumps independently controlled by proprietary Dolomite software. The mineral oil continuous phase was controlled by a syringe pump. The fluids were fed into the droplet chip through 4 equal lengths of polytetrafluoroethylene (PTFE) microbore tubing (Figure 1).

**Figure 1.**
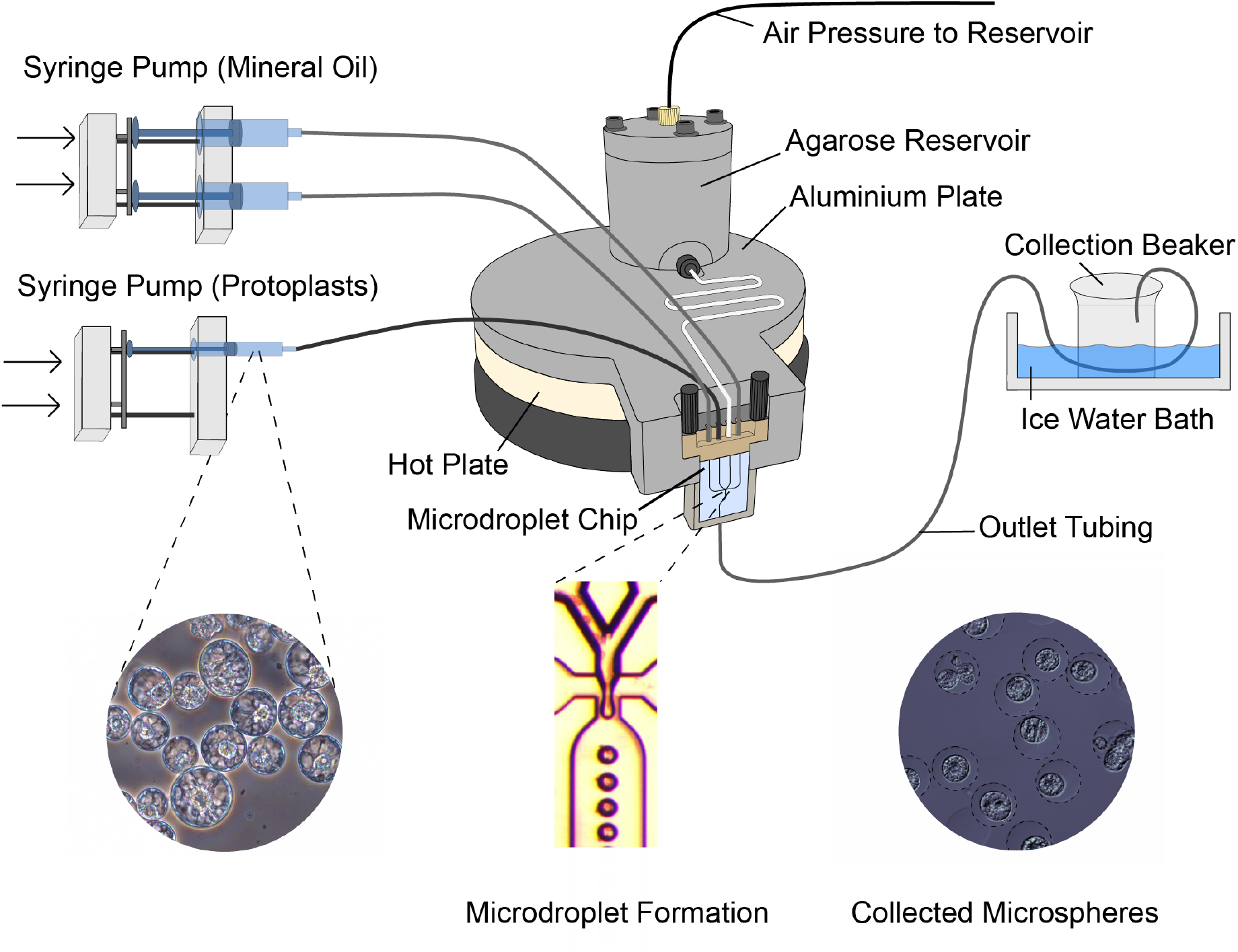
Agarose microsphere encapsulation system. Two separate syringe pumps control the flow of protoplasts and mineral oil into a Dolomite Microfluidics 4-channel, 2-reagent, hydrophobic microdroplet chip (#3200725). Agarose flows from a pressurized and heated reservoir into the microdroplet chip, with input line temperatures maintained at 35×C by a channeled aluminum plate and liquid agarose reservoir atop a digital hot plate. Outlet tubing carries liquid microdroplets in oil through an ice water bath that cools the agarose into solid microspheres before depositing them into a sterile collection beaker. Agarose microspheres taken from the collection beaker are washed and centrifuged.

Of the two central aqueous channels one was reserved for the protoplasts prepared in Step 1. The second central aqueous channel was fed from a warmed reservoir containing 1.5% low-gel-temperature agarose (Sigma-Aldrich) dissolved into PWS #1 and maintained at a temperature of 35×C. Agarose solution was prefiltered through 0.22 µm syringe filters for sterilization.

With appropriate flow control, a stream of monodisperse agarose droplets was formed at the chip junction where the aqueous and oil phases intersect (Figure 1). Droplet diameter can be increased or decreased by adjusting the flow rates of the continuous and discontinuous phases. Liquid droplets exit the chip into outlet tubing which carries them through an ice bath causing microdroplets to solidify into microspheres before being deposited into a collection beaker. (Figure 2).

**Figure 2.**
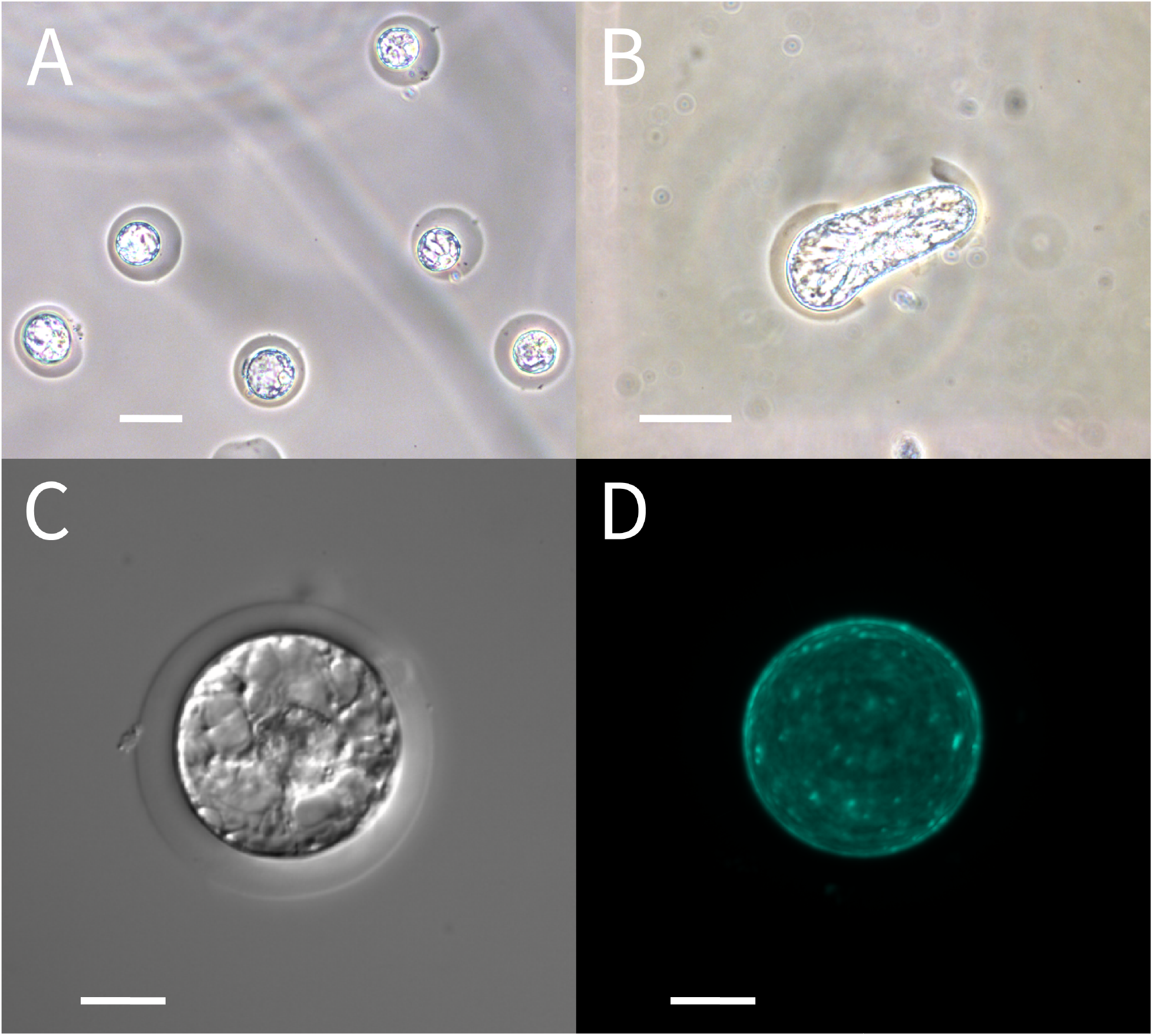
(A) Phase-contrast image of BY-2 protoplasts in solidified microspheres suspended in MS medium immediately after encapsulation. (B) Phase-contrast image of a BY-2 cell rupturing the agarose microbead 6 days after encapsulation. (C) Nomarski differential interference contrast image of an individual BY-2 Cell 24 h after encapsulation showing active cytoplasmic streaming. (D) Calcofluor White fluorescence of the cell in Fig. 2C showing regeneration of a thin cell wall. Scale bars = 50 μm (2A, 2B) and 20 μm (2C, 2D) Figure adapted from (Grasso & Lintilhac, 2016).

Consistently sized (ca 60µm) spherical hydrogel microspheres were successfully generated at a rate of approximately 130 droplets/sec. Individual protoplasts were successfully encapsulated with good viability as determined by cytoplasmic streaming activity shortly after encapsulation (Figure 2C). Cellulose staining with Calcofluor White showed that 24 hours after initial protoplast release all living cells had regenerated a thin surrounding wall (Figure 2D). The amount of cell wall cell wall regenerated continued to increase up to 6 days after encapsulation in agarose microspheres. Living cells proceeded to elongate and divide, eventually bursting the agarose microsphere in which they had originally been encapsulated (Figure 2B). Immediately following droplet formation, approximately 25% of droplets contained protoplasts. (Grasso and Lintilhac 2016).

Agarose encapsulated protoplasts in mineral oil were transferred from the collection beaker to a 15 mL conical centrifuge tube and 6mL of PWS #1 was added. The tube was centrifuged for 4 minutes at 800 RPM forming a pellet of agarose microspheres as they moved down from the oil layer into the aqueous phase. The bottom of the tube was punctured with a sterile needle and microspheres were drained out and collected free of oil. This solution was filtered through a sterilized 149-micron nylon mesh to remove large pieces of debris. At this point microspheres were centrifuged again to form a pellet and then transferred to PCM. They were placed in an incubator set to 27×C with no shaking.

### Step 3

Dual-layer agarose and alginate-methacrylate microbead encapsulation of agarose microspheres (Figure 3).

**Figure 3.**
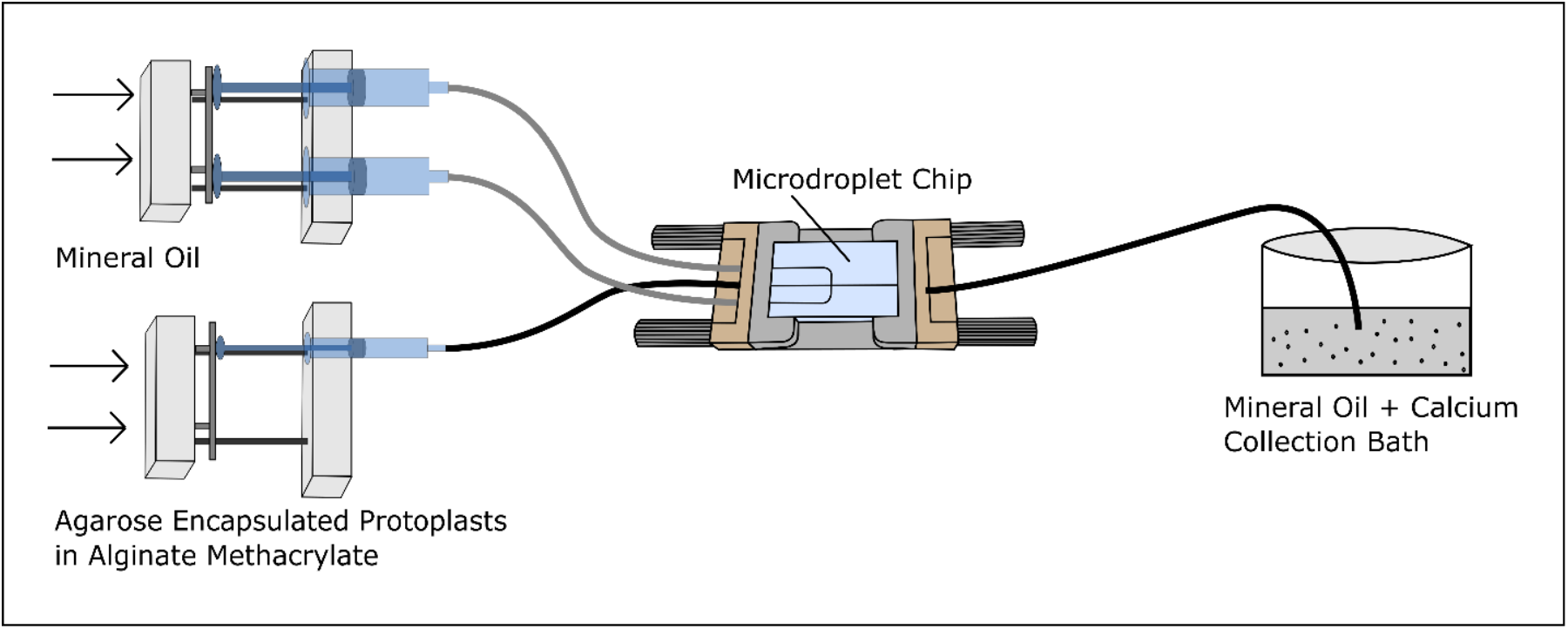
Agarose encapsulated protoplasts are pre-mixed into an AlgMA stream flowing into faster moving mineral oil resulting in agarose microspheres encased in a layer of AlgMA.

Agarose encapsulated protoplast microspheres were collected after 24 hours of culture and centrifuged at 100g in a 15mL conical centrifuge tube. After the supernatant was decanted microspheres were re-suspended in protoplast washing solution PWS #2 and allowed to rest for 15 minutes to wash out any salts found in the PCM that might cause premature gelling of the alginate-methacrylate (AlgMA). The washed microspheres were centrifuged to form a pellet and re-suspended in PWS w/o MgCl_2_ to a final density of approximately 650 agarose microspheres/µl.

AlgMA solution was prepared by first sterilizing 0.02g of dry AlgMA polymer by saturating it with 100% ethyl alcohol and allowing the alcohol to evaporate in a sterile cabinet. Once evaporated, the dry sterile polymer was dissolved in 0.7mL of AMDS by shuttling the solution and polymer between two 3mL syringes connected by a Luer couple for 30 minutes. Once fully dissolved, 300µl of agarose microsphere encapsulated protoplasts in PWS w/o MgCl_2_ were added to the AlgMA solution and mixed by slowly shuttling back and forth between the two syringes ~30 times. Encapsulation within agarose protects protoplasts from shearing forces caused during Step 3 which results in agarose microspheres being suspended in a liquid (ungelled) AlgMA carrier in anticipation of a final round of microfluidic droplet encapsulation.

### Step 4

Dual layer AlgMA microbead encapsulation protocol:

AlgMA microbead generation was done using a 3-channel, 190µm junction diameter, hydrophobic, flow-focusing microdroplet chip (# 3000437) from Dolomite Microfluidics (Figure 4). The continuous phase was mineral oil with 2% Span 80. The flows of the AlgMA solution and mineral oil were controlled separately by two syringe pumps (Figures 3 and 4).

**Figure 4.**
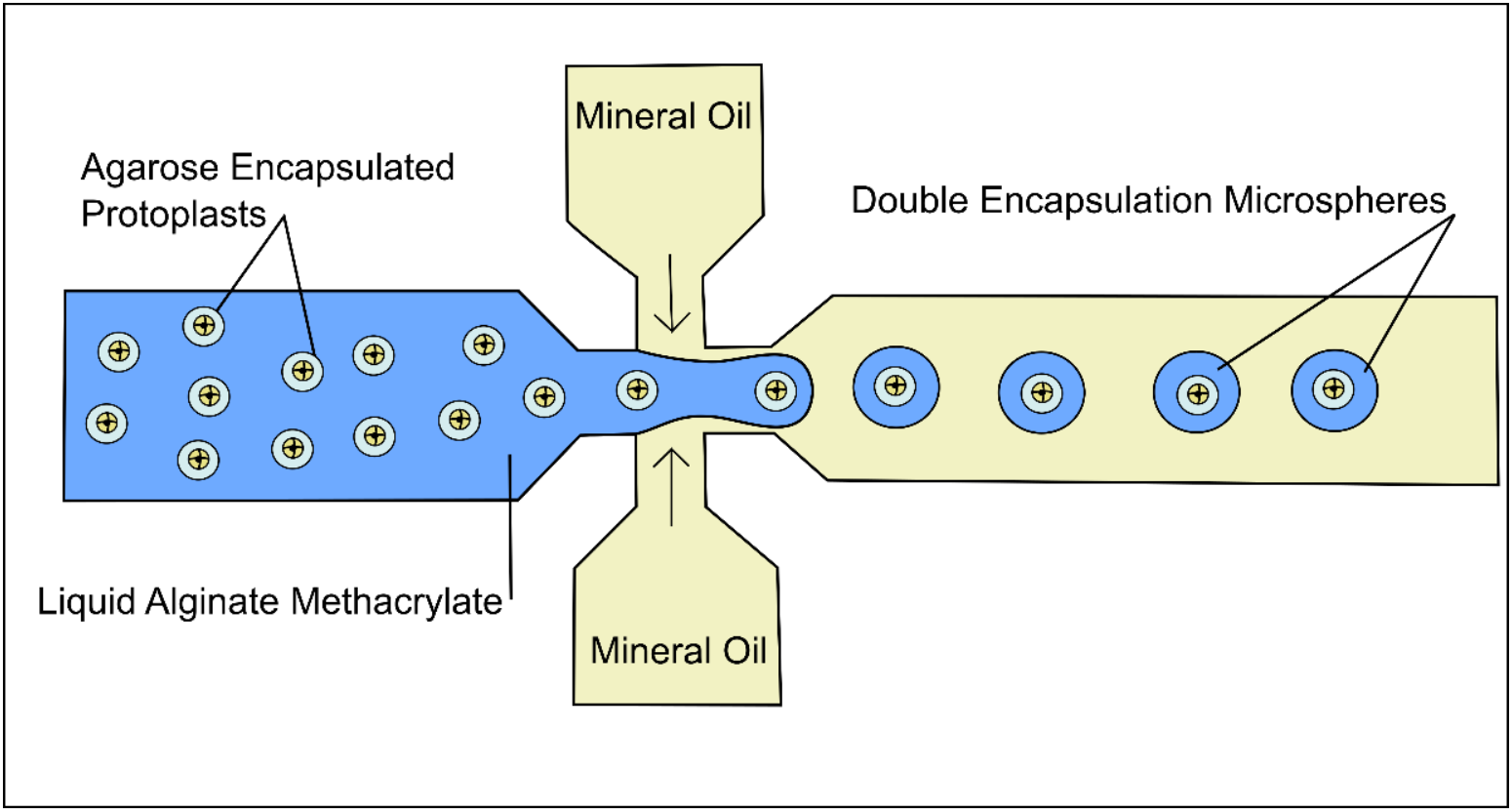
Schematic of the 3-channel microdroplet chip used for AlgMA double encapsulation.

### Step 5

final collection and gelation of microbeads.

For the final collection of microbeads, dry CaCl_2_ was added in powder form to 5mL of mineral oil and 2% span 80 (10mM Ca^2+^ equivalent) as an initial cross-linking agent and mixed with a magnetic stir bar for 2 hours before final encapsulation. During encapsulation the CaCl_2_ mineral oil bath was continuously stirred (Figure 4). A schematic of the AlgMA encapsulation system is shown in Figure 4. The syringe pump controlling the flow of oil was set to 1000 µl/hour and the syringe pump controlling the flow of protoplasts in AlgMA was set to 150 µl/hour. The system was run for one hour. The collected calcium crosslinked AlgMA microbeads in mineral oil (Figure 5) were added to a 15mL conical centrifuge tube along with 6mL Calcium Solution (CS) and centrifuged at 100 RPM for 4 minutes to transfer microbeads from the mineral oil layer to the CS. AlgMA microbeads were drained from the bottom of the tube and allowed to sit in the CS for 10 minutes to ensure complete calcium crosslinking. After 10 minutes microspheres were pelleted again by centrifugation and the CS supernatant was removed and replaced with 3mL of PCM. Dual layer microspheres were transferred to a small petri dish and placed in an incubator at 27×C. Note that there are two stages to the Calcium crosslinking, the first using powdered CaCl_2_ in oil, and the second using CaCl_2_ in aqueous solution.

**Figure 5.**
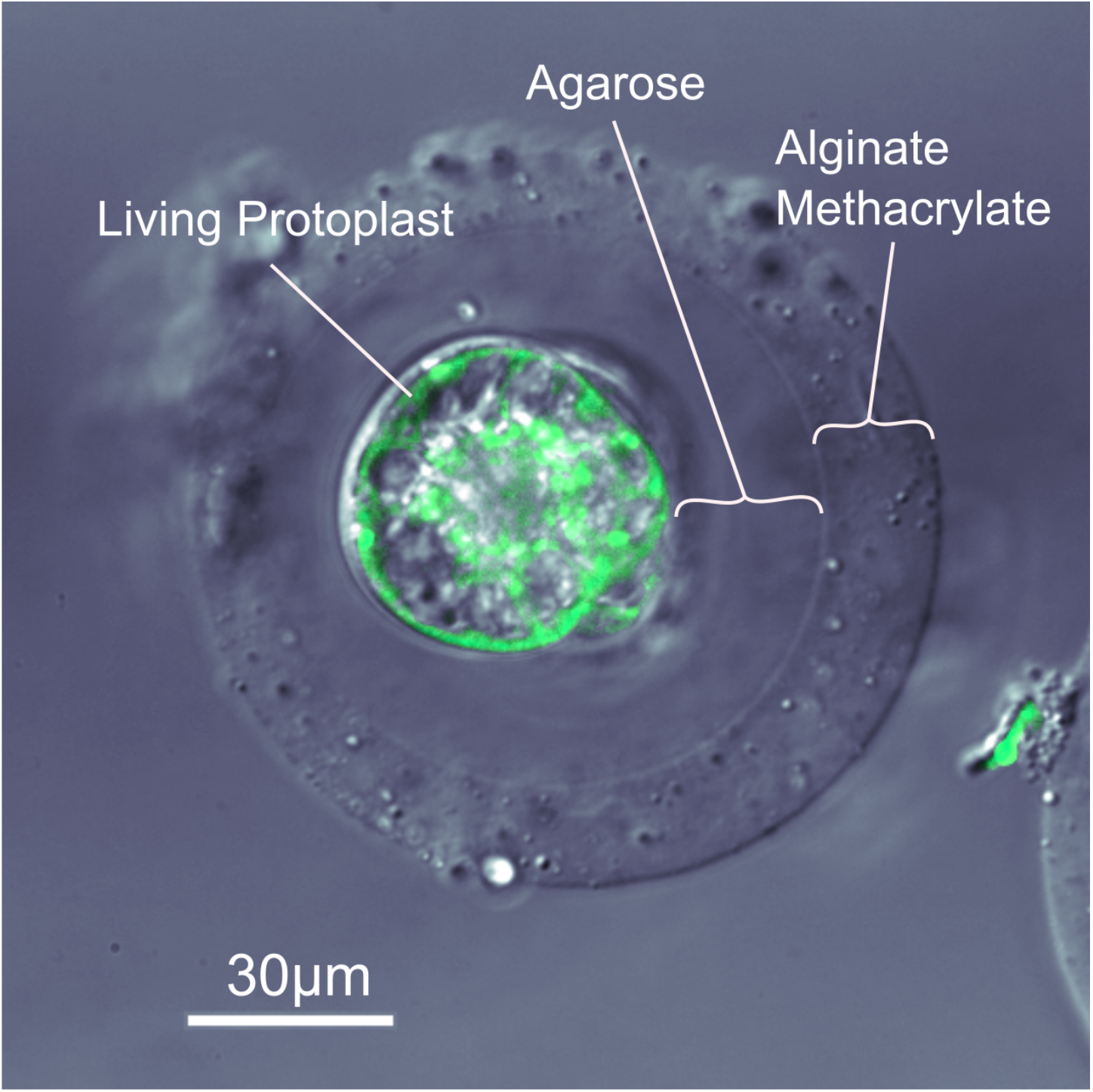
Overlay of a differential interference contrast and fluorescence image of an AlgMA/ agarose dual-layer microbead encapsulating a living protoplast. Image was taken 3 days after protoplast isolation and 2 days after AlgMA encapsulation. The living protoplast is stained green with FDA viability stain. Labels show the outer layer of AlgMA and the inner layer of agarose that encapsulates the living protoplast.

**Figure 6.**
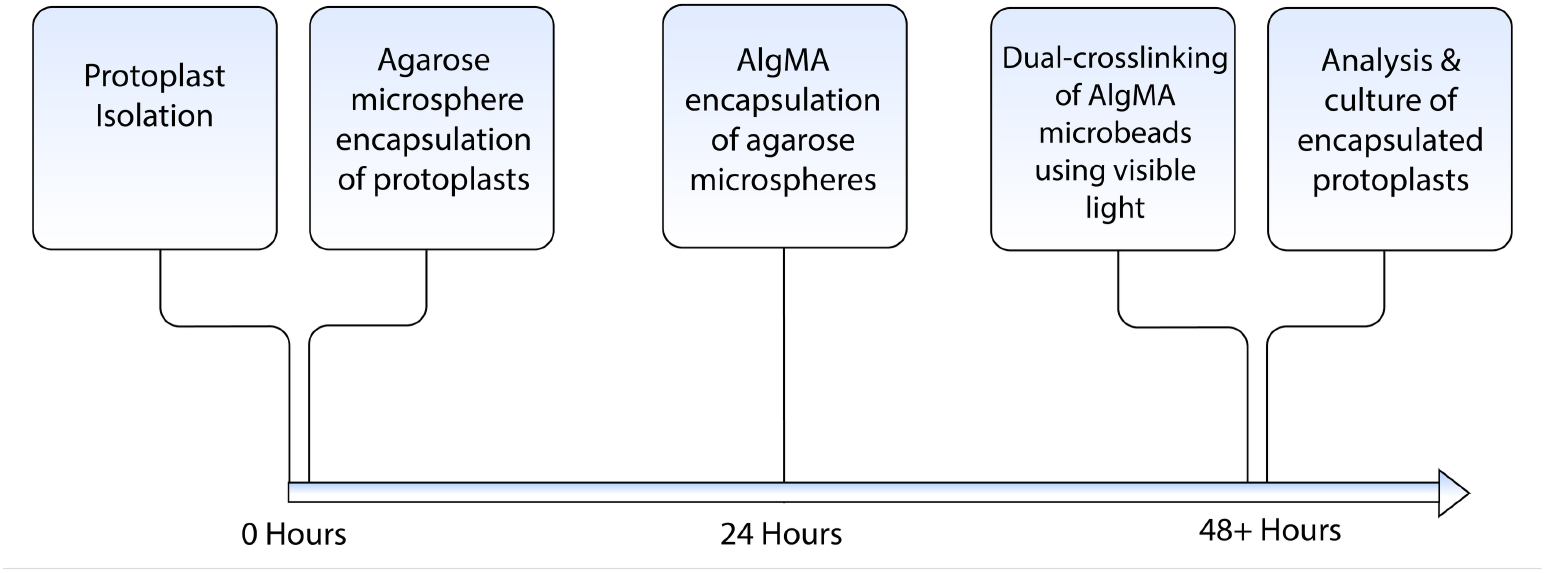
Live protoplast double-encapsulation timeline.

### Buoyancy control

Precise control of protoplast buoyancy in the suspending medium is essential. Freshly prepared protoplasts, being denser than the suspending medium will settle onto the lower, inner surface of the microbore feed lines where they will tend to clump. It is essential to keep them uniformly suspended in the carrier medium. There are two ways to control protoplast buoyancy. Protoplast density can be controlled osmotically by adjusting the osmolarity of the suspending medium, making it possible to adjust protoplast densities by causing them to shrink or swell. We use freezing-point osmometry to follow medium osmolarity at every step. A second approach to buoyancy control is to adjust the specific gravity of the medium by introducing a high specific-gravity additive in calibrated amounts, adjusting the density of the medium to match that of the protoplasts. We have found that a high-density, non-toxic radiological contrast agent such as Iodixanol (Fisher OptiPrep™) can be used to maintain neutral density at critical times during microdroplet and microbead assembly, as well as controlling buoyancy during centrifugal phase separation. Iodixanol is non-toxic and does not significantly alter osmotic balance with changing concentration. It can be pH adjusted, and autoclave sterilized without disrupting its properties.

## Results

We were able to generate dual layer AlgMA microbeads in small numbers, losing a proportion of viable protoplasts at each step along the way. However, we have successfully demonstrated that it is possible to generate multilayered microbeads encapsulating living plant cells that remained viable for 3 days (Figure 5).

Single layer protoplast/agarose microspheres were premixed into prepared AlgMA solution. Dual layer AlgMA microbeads (Figure 5) were generated using a flow-focusing microfluidic chip and a continuous phase consisting of mineral oil with 2% Span 80. The calcium-oil collection bath crosslinked AlgMA droplets into consistently shaped microbeads. The crosslinked AlgMA microbeads were separated from the calcium-oil collection bath by centrifugation into aqueous medium. While viability of dual layer microbead/protoplasts was low, staining with fluorescein diacetate revealed living plant cells (Figure 5). Total elapsed time from initial protoplast release to microbead assessment was 3 days.

Previous work has shown that viable single-layer encapsulation of living plant protoplasts is possible (Grasso and Lintilhac 2016). We can now demonstrate that multi-layer encapsulation generating spherical dual-layer droplets, each containing a viable BY-2 protoplast is also possible. In addition, we show that dual-layer microbeads can be formed using different polymer systems with distinct mechanical behaviors, and that the dimensions of the resulting microbeads are within the range of native sporangial structures.

### Unresolved issues

In recent work we have outlined the underlying biomechanical logic of archesporial differentiation in the earliest stages of reproductive differentiation in the land plants (Lintilhac, 2024). Here we address the question of how the mechanical stimuli that target precursors of the archesporium can be approached experimentally. Our work does not address the question of sourcing different cell lines from intact plant tissues, nor does it attempt to address the question of developmental competence which may be established independently through hormonal pathways (Di Mambro, Et Al, 2017).

## Discussion

The work reported here is a preliminary study. We believe that with further development it may lead to a reliable way to manipulate the mechanical environment of individual plant cells, potentially adding a new suite of experimental tools to the study of plant biomechanics at the single cell level. A possible future outcome of this work would be the ability to elicit the differentiation of supernumerary archesporia *in vitro* and the subsequent induction of meiosis using engineered biomimetic sporangia which can be produced in large numbers, leading to the possibility of harvesting haploid pseudo-gametes for various purposes.

The list of experimental tools available for tuning the mechanical properties of hydrogel materials is growing. Further development of the methacrylated alginate methodology described here may allow the outer layer to be photo-crosslinked by free radical polymerization, making the mechanical properties of the microspheres subject to modification post-encapsulation (Etter et al. 2018). This would also result in covalent bonds between alginate chains which would then be protected from destabilization in physiological solutions like the ionic bonds present in calcium crosslinked alginate. Future advances in hydrogel engineering and cross-linking methods (Tahir and Floreani, 2022) using mechanically active hydrogel polymers capable of controlled shrinking or swelling (Takashima et al 2012) (Pan et al 2016)) (Sugaya et al, 2013) (Yang, et al 2013) raise the possibility of actively manipulating the intensity and location of center-focused stress fields (isotropic points) acting on appropriately competent living plant cells at targeted locations, under conditions that mimic those of living plant sporangia.

Although genetically scripted molecular controls necessarily underlie all developmental processes, there are intrinsic limits to the ability of molecular signals to coordinate precise structural decisions at the transcellular level. Information flows based on molecular population changes are necessarily stochastic at the tissue level (Lintilhac 2022), whereas physical signals, being intrinsically deterministic and therefore less prone to stochastic dissipation, do not rely on molecular transport at all, and can be directed to carry out action at a distance instantly.

The production of physical shape and form in plants necessarily reflects the material properties of their structural components and the way they interact with the mechanical signals imposed by the surrounding cellular environment; and while it is true that the details of the mechanical stress environment and surface topography can be tuned at the molecular level by transcriptionally directed inputs, the deterministic physical interactions that drive architecture at the cellular level must be understood independently. In turgid and growing plant organs, including nascent sporangial structures, the physical forces released into the body of the growing sporangium reflect the growth potential of the constituent cells, but they also reflect the geometry and surface topography of the structure as a whole, providing a continuously changing landscape of structure-specific physical information which is seamlessly updated with the changing shape of the sporangium as it continues to grow.

## Conclusion

Physical and mechanical inputs are increasingly recognized as pivotal in the development of plant form and function (Khadka et al, 2020), and while the role of cell and tissue mechanics is correctly seen against a background of molecular control systems, we are beginning to understand that in the plant kingdom, developmental systems have evolved within the constraints of apoplastic continuity, immotile cells, and pressure-driven growth. Under these constraints cell and tissue mechanics have evolved into robust informational networks that can operate in environments where stochastic molecular systems cannot.

PCT/US2023/027943 patent pending

